# Spatial and temporal control of 3D hydrogel viscoelasticity through phototuning

**DOI:** 10.1101/2023.08.07.551544

**Authors:** Philip Crandell, Ryan Stowers

## Abstract

The mechanical properties of the extracellular environment can regulate a variety of cellular functions, such as spreading, migration, proliferation, and even differentiation and phenotypic determination. Much effort has been directed at understanding the effects of the extracellular matrix (ECM) elastic modulus and more recently, stress relaxation, on cellular processes. In physiological contexts like development, wound healing, and fibrotic disease progression, ECM mechanical properties change substantially over time or space. Dynamically tunable hydrogel platforms have been developed to spatiotemporally modulate a gel’s elastic modulus. However, dynamically altering the stress relaxation rate of a hydrogel remains a challenge. Here, we present a strategy to tune hydrogel stress relaxation rates in time or space using a light-triggered tethering of poly(ethylene glycol) (PEG) to alginate. We show that stress relaxation rate can be tuned without altering the elastic modulus of the hydrogel. We found that cells are capable of sensing and responding to dynamic stress relaxation rate changes, both morphologically and through differences in proliferation rates. We also exploited the light-based technique to generate spatial patterns of stress relaxation rates in 3D hydrogels. We anticipate that user-directed control of 3D hydrogel stress relaxation rate will be a powerful tool that enables studies that mimic dynamic ECM contexts, or as a means to guide cell fate in space and time for tissue engineering applications.

## INTRODUCTION

Hydrogels are widely used to simulate aspects of the native ECM for in vitro cell culture^1^. Key biophysical and biochemical properties of the ECM can be precisely controlled in hydrogel cultures, enabling reductionist investigations into the effect of the cellular microenvironment or design of artificial ECMs for in vitro cell and tissue models^2^. The mechanical properties of the hydrogel network are particularly important in regulating cell behaviors, such as cell proliferation, adhesion, migration, and spreading. Indeed, the stiffness (elastic modulus) of the hydrogel can direct differentiation of stem cells, enhance cancer cell malignant traits, enable or restrict vascularization and control organoid formation^3–8^. While the importance of mimicking or controlling the elastic modulus of a hydrogel matrix is now well-recognized, most soft tissues in the body do not behave purely elastically, but instead exhibit properties of both elastic solids and viscous fluids^9,10^. The viscous component of the tissues dissipate energy over time, resulting in time-dependent mechanical phenomena, such as stress relaxation – a decrease in stress under a constant strain, or creep – an increase in strain under constant stress. Recently, differences in matrix viscoelasticity have been shown to impact cell fate and phenotype as well, underscoring the importance of mimicking this mechanical behavior in in vitro models of the cellular microenvironment^11–16^.

Many widely used hydrogel crosslinking chemistries form irreversible covalent bonds and the resulting hydrogels behave elastically; that is, stress is stored indefinitely upon deformation, not dissipated over time. In order to generate highly viscoelastic gels, the polymer network must be able to be rearranged in response to force through disruption of the network crosslinks. Several strategies have been employed towards this end, including the use of ionic crosslinks, guest-host or other supramolecular interactions, dynamic covalent bonds, and hydrophobic interactions^11,13,17,18^. These hydrogel systems have been used to interrogate the cellular response to viscoelasticity, a burgeoning area in mechanobiology and 3D culture models. Mesenchymal stem cell spreading, focal adhesion formation and osteogenic differentiation is significantly enhanced in fast relaxing matrices^11–14^. Viscoelastic matrices have been used to generate, expand and passage stem cell-derived organoids, and the extent of local viscoelasticity can enable morphogenetic processes such as intestinal organoid budding^16,19–21^. The tumor microenvironment is also viscoelastic, and cancer cells are responsive to different rates of stress relaxation through different rates of proliferation, migration, and modes of invasion^22–25^.

Tissue mechanical properties vary throughout time during many biological processes such as development and aging, and diseases such as organ fibrosis and solid tumor progression^26–30^. Similarly, tissues are spatially heterogeneous in terms of mechanical properties^9,31^. Dynamically tunable hydrogels have been developed to mimic the spatial and temporal changes that occur in tissues^32–40^. These approaches primarily rely on methods to alter hydrogel crosslink density in a temporal or spatial gradient, or with a user-directed stimulus like light, heat, or ultrasound. Crosslink density is directly related to the hydrogel network elasticity; thus the stiffness of these gels can be altered spatiotemporally. However, altering the viscoelasticity of a hydrogel in time or space, for example the rate at which stress relaxes under constant strain, is more challenging. There are two recent examples using PEG gels in which a light-triggered exchange of crosslink bonds is utilized to permit stress relaxation, enabling control of viscoelasticity^21,41^. Here, we demonstrate a strategy to spatiotemporally modulate 3D hydrogel stress relaxation without altering stiffness and in the presence of cells. Our platform utilizes alginate gels crosslinked with calcium, which are inherently viscoelastic due to the nature of the ionic bonds. Modification of alginate with monofunctional PEG chains can enhance the stress relaxation rate of the gels in a concentration-dependent manner^42^. Our approach is to employ photoclick chemistry to conjugate PEG chains to an existing, 3D, cell-laden alginate network to modify its viscoelasticity without altering the stiffness of the gel (Fig. 1).

**Figure 1:**
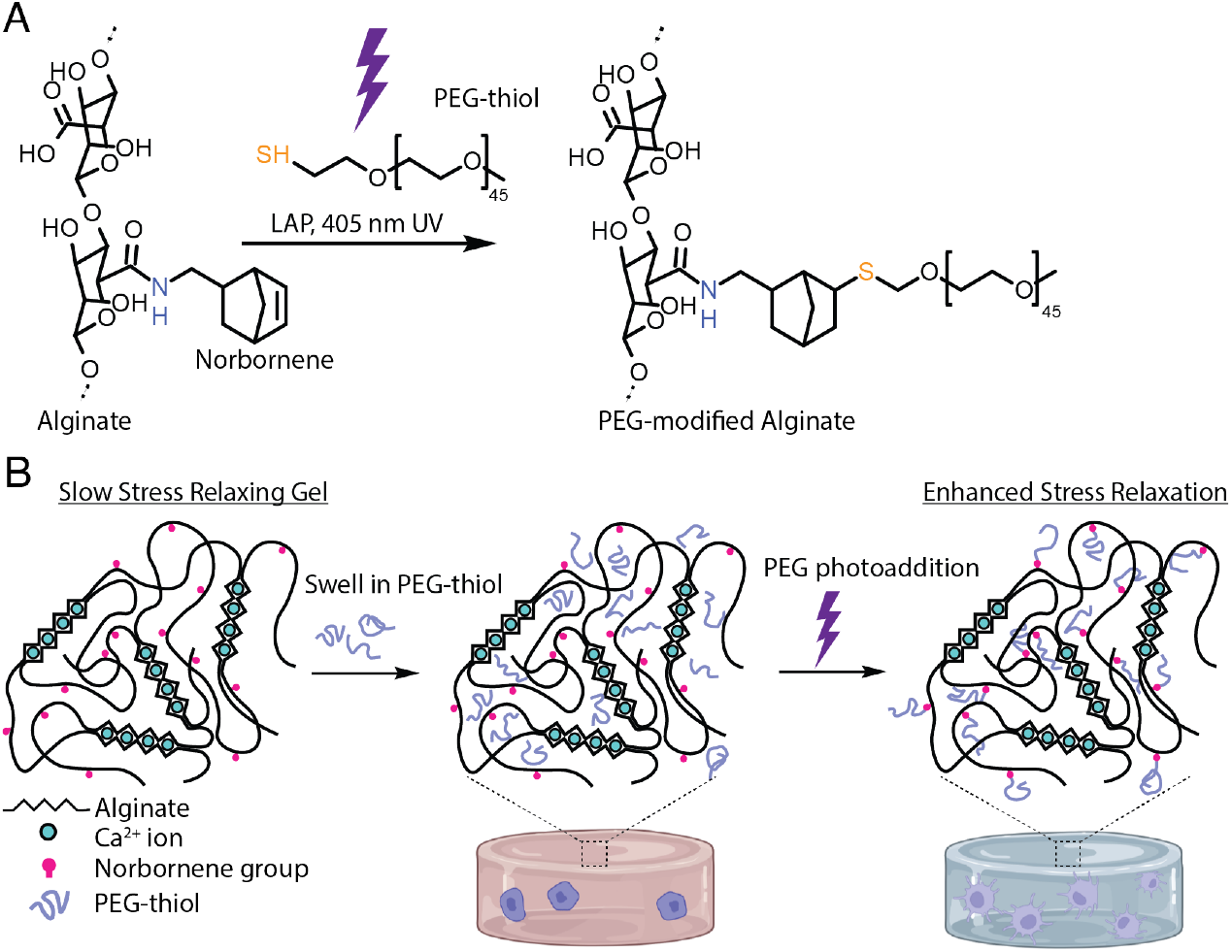
Strategy to generate light-triggered changes in alginate hydrogel stress relaxation rate. (a) Chemical structures of alginate modified with norbornene and thiol-ene reaction to conjugate PEG to alginate in the presence of 405 nm light and the photoinitiator lithium phenyl-2,4,6-trimethylbenzoylphosphinate (LAP). (b) Schematic of alginate polymer network during photoaddition of PEG chains that serve to enhance the stress relaxation rate of the gel.

## MATERIALS AND METHODS

### Alginate preparation

Alginate (280 kDa molecular weight, LF20/40) from FMC biopolymer was dissolved at 1% in deionized water and dialyzed with 10 kDa MWCO membranes against deionized water for 3 days. Following dialysis, alginate was purified with activated charcoal, sterile filtered, frozen, and lyophilized.

### Functionalization of alginate with RGD

RGD peptides were coupled to alginate using carbodiimide chemistry. Alginate was dissolved in 0.1 M 2-(N-morpholino)ethanesulfonic acid (MES) buffer of pH 6.5. Then, appropriate amounts of sulfo-NHS (N-hydroxysulfosuccinimide), N-(3-dimethylaminopropyl)-N′-ethyl carbodiimide hydrochloride (EDC), and the peptide sequence GGGGRGDSP were mixed and the reaction was left to stir for 20 h at room temperature^**14,43**^. The product was then transferred to 10 kDa MWCO dialysis tubing and dialyzed against decreasing concentration NaCl solutions starting from 120 mM to 0 mM over 2 days followed by 1 day of dialysis against deionized water. Water was then removed via lyophilization to yield functionalized alginate.

### Functionalization of alginate with norbornene

Alginate functionalized with RGD was additionally functionalized with norbornene using a similar procedure to the above. Alginate-RGD was dissolved in MES buffer and functionalized with sulfo-NHS, EDC, and norbornene (5-norbornene-2-methylamine, TCI Chemicals). After allowing sulfo-NHS, EDC and alginate-RGD to dissolve, the pH of the MES was raised to 8, and norbornene was added. Following the reaction, the alginate was dialyzed and lyophilized as described above. Lyophilized alginate was dissolved in phenol red-free Dulbecco’s Modified Eagle Medium (DMEM) at 3% M/V. Free norbornenes were quantified by reacting with 2 kDa mPEG-thiols and Ellman’s reagent (5,5-dithio-bis-(2-nitrobenzoic acid)). The remaining thiols were quantified with Ellman’s reagent to determine norbornene substitution of alginate.

### Hydrogel formation and tuning of mechanical properties

To tune the viscoelasticity of alginate after gelation, 2 kDa mPEG-thiol chains were reacted with norbornene groups on alginate. Alginate gels were formed, as described previously^**44**^. Briefly, a syringe containing alginate was coupled to a second syringe containing calcium sulfate in DMEM and the contents of both syringes were rapidly mixed. Hydrogels were cast directly into 8-well chambered cover glasses or cast between two silanized glass plates spaced 2 mm apart and punched into 8 mm diameter discs. After gelation for 40 minutes at 37oc, gels were equilibrated in DMEM. For photocoupling, PEG-thiol and LAP (lithium phenyl-2,4,6-trimethylbenzoylphosphinate) in phenol red-free DMEM were added to the well either after immediately post-gelation, or after 24 or 72 hours. PEG was allowed to swell in for 4 hours, then gels were exposed to 60 seconds of 405 nm light, and the media was changed to remove any unreacted PEG.

### Mechanical characterization

Hydrogel mechanical properties were characterized on a TA Instruments ARES G2 strain-controlled rheometer with 8 mm parallel plates. Alginate hydrogels with and without PEG were formed as described above. All hydrogel samples were measured 24 hours after photoaddition of PEG, if applicable. The top plate was brought down until it registered a non-negative axial force, and the gap between plates was filled with DMEM. Shear modulus was measured using a frequency sweep from 0.1 to 10 Hz at a strain of 1%. The elastic modulus (E) was calculated assuming a Poisson’s ratio (ν) of 0.5 using the equation:

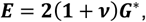

where the complex modulus (G*) was determined from the storage modulus (G’) and loss modulus (G’’) using:

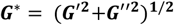

For stress-relaxation tests, a constant strain of 15% was applied, and stress was recorded over time. Relaxation time was defined as the time taken for the stress to relax to half of its initial value. For creep tests, a constant 100 Pa stress was applied to each alginate gel for 3600 s and the gel was allowed to recover at 0 Pa applied stress for 7200 s. Control values for each creep-recovery test were derived via a frequency sweep performed directly before the test.

### Cell culture

Mouse D1 MSCs (ATCC, CRL-12424) were cultured in DMEM with 10% fetal bovine serum and 1% penicillin/streptomycin. Mouse D1 cells were passaged at approximately 50% confluency and the media was changed every 48 hours. MDA-MB-231 cells (ATCC, HTB-26) were expanded in DMEM containing 10% fetal bovine serum and 1% penicillin/streptomycin. The medium was changed every 48 hours and the cells were passaged at 70% confluency.

### Staining and microscopy

To evaluate cell morphology, cells were cultured in hydrogels cast into a chambered cover glass (Nunc Lab-Tek II). After the last day of the culture period, cells were fixed using 4% paraformaldehyde in serum-free DMEM at 37°C for 1 hour. Gels were then washed 3 times in PBS containing calcium and 0.1% Triton X-100 for 30 minutes each. Alexa Fluor 555 phalloidin and DAPI were added for 90 minutes at room temperature, and the sample was again washed 3 times with calcium PBS at 30-minute intervals. Samples were imaged immediately using a Leica SP8 laser scanning confocal microscope with a 25x magnification water immersion objective.

### Proliferation assay

To quantify proliferation, cells were encapsulated in hydrogels as described above and media containing 10 μM EdU (5-Ethynyl-2’-deoxyuridine) was added 24 hours before fixation. Gels were fixed in 4% paraformaldehyde for 45 minutes, washed 3 times with PBS containing calcium, and incubated with 30% w/v sucrose in calcium-containing PBS overnight. Gels were then placed in a mixture of 50% sucrose and 50% Tissue Tek O.C.T. compound on a shaker for 8 hours before being frozen in O.C.T. and sectioned. Sectioned gels were stained for EdU using a 647 fluorescent EdU kit (Click-&-Go EdU 647, Click Chemistry tools) per the manufacturer’s directions. After functionalizing EdU with fluorophore, sections were incubated in 1:1000 DAPI for 30 minutes and washed 3x with PBS.

### Photopatterning of alginate

Patterned photomasks were produced using a laser printer to transfer toner to an 8×10 inch polystyrene sheet (Shrinky Dink). Sheets were then cut and placed in an oven at 160°C for two minutes, similar, to previous methods^**45**^. A glass slide was placed on the sheet as it shrunk to ensure the pattern remained flat. The patterns were transferred to the gels in a chambered coverglass using a collimated 405 nm laser (NDV4512, Laserlands) by placing the photomask against the glass surface of the sample and illuminating through it for 30 seconds.

### Image analysis

All images were collected using a Leica SP8 confocal microscope using a 0.95 NA 25x magnification water immersion objective. Metrics describing cell morphology in three dimensions such as sphericity and volume were quantified using Bitplane Imaris 9.5 software. In Imaris, sphericity is defined as the ratio of the surface area of a sphere with the same volume as the cell to the surface area of the cell itself. Solidity of a maximum projection of a 3D stack was quantified using ImageJ, where solidity represented the difference between the convex hull area and the area of the cell itself. All 2D images of cell proliferation staining were analyzed and quantified by counting the number of co-stained DAPI and EdU cells and taking that as a fraction out of each separate field of view. Analysis of morphology in patterned gels was performed from a single stitched stack from 3 technical replicates, each 50 μm deep and 3 mm^2^ area.

### Statistical analysis

Statistical comparisons were performed using GraphPad Prism 9.5. One-way analysis of variance was used to compare more than two groups. For measurements like cell volume, sphericity, and solidity the D’Agostino-Pearson normality test was first performed to test if the data could be treated normally. For cell morphology experiments approximately 25 cells were collected per trial and data from 3 separate trials was pooled for analysis. A total of 18 fields of view were analyzed from 2 separate trials in each condition. Image analysis of patterned gels was performed from a single stitched stack from 3 technical replicates. Cells from photopatterned gels were analyzed using 2D metrics because the vertical sampling rate was insufficient for 3D analysis, and values for circularity, roundness, and solidity were reported.

## RESULTS AND DISCUSSION

### Photocoupling of PEG to 3D alginate hydrogel networks enhances stress relaxation rate

First, we modified high molecular weight alginate, previously shown to have slow stress relaxation, with norbornene functional groups to enable photoclick conjugation of monofunctional PEG-thiol chains^**11**^. Norbornene methylamine was grafted to alginate using carbodiimide chemistry, a well-established chemistry^**46**^ for functionalizing amines to alginate’s carboxylic acid groups (Fig. 1A). Quantification of this reaction using Ellman’s reagent found that 7% of the carboxylic acids were substituted with free thiols. Norbornene functional groups can react with free thiols in solution in the presence of a photo-initiator via the cytocompatible thiol-ene reaction (Fig. 1A). Alginate modified with PEG chains before gelation has been shown to have increased stress relaxation compared to unmodified alginate, and the stress relaxation rate increases with the amount of PEG added^**44,47**^. Here, we demonstrate the extent to which stress relaxation can be tuned by photocoupling of PEGs in a pre-formed 3D gel and in the presence of cells (Fig. 1B).

To characterize the mechanical changes in alginate gels after PEG-addition in conditions that mimic cell culture situations, norbornene-alginate hydrogels were ionically crosslinked, and cut into 8 mm cylindrical samples with a biopsy punch to ensure samples had the same initial mechanical properties. To modify mechanical properties after gelation, gels were equilibrated with varying concentrations of monofunctional 2 kDa PEG-thiol and the photoinitator LAP, exposed to a 405 nm laser, then swollen in buffer to remove any unreacted PEG chains (Fig. 2A). Alginate hydrogels with more photocoupled PEG, presented as the ratio of PEG-thiols to norbornenes, produced faster relaxing hydrogels, demonstrating proof-of-concept of our approach (Fig. 2B,C). For ease of comparison of stress relaxation rates, we determined the time necessary to reach half of the initial stress (τ_1/2_). Relaxation half-times varied from 840 seconds for unmodified alginate to 82 seconds for hydrogels where the concentration of added thiol exceeded the number for norbornene groups. Adding PEG in a 1.6 molar excess to norbornene produced similar stress relaxation times to a 1.2 molar excess, indicating that there is a diminishing effect at high PEG concentrations. The elastic moduli were not significantly affected by PEG photoaddition, demonstrating that these two mechanical parameters can be independently modulated (Fig. 2D). Creep-recovery tests were performed on unmodified and 1.2:1 thiol: norbornene hydrogels. In a creep-recovery test, strain is measured while a constant stress is applied to the gel (creep), followed by a period of zero stress (recovery). PEG-modified hydrogels had significantly higher residual strains after creep-recovery testing, indicating more plastic deformation in these gels (Fig. 2E). Together these results show that photoaddition of PEGs in alginate hydrogels can allow for on-demand changes in viscoelastic behavior of alginate hydrogels independent of the elastic modulus. Importantly, this range of stress relaxation rate, from approximately 100 to 1000 seconds, spans the measured relaxation rates for many soft tissues^**10**^ and the spectrum over which cellular behaviors are drastically altered^**11,14,47**^.

**Figure 2:**
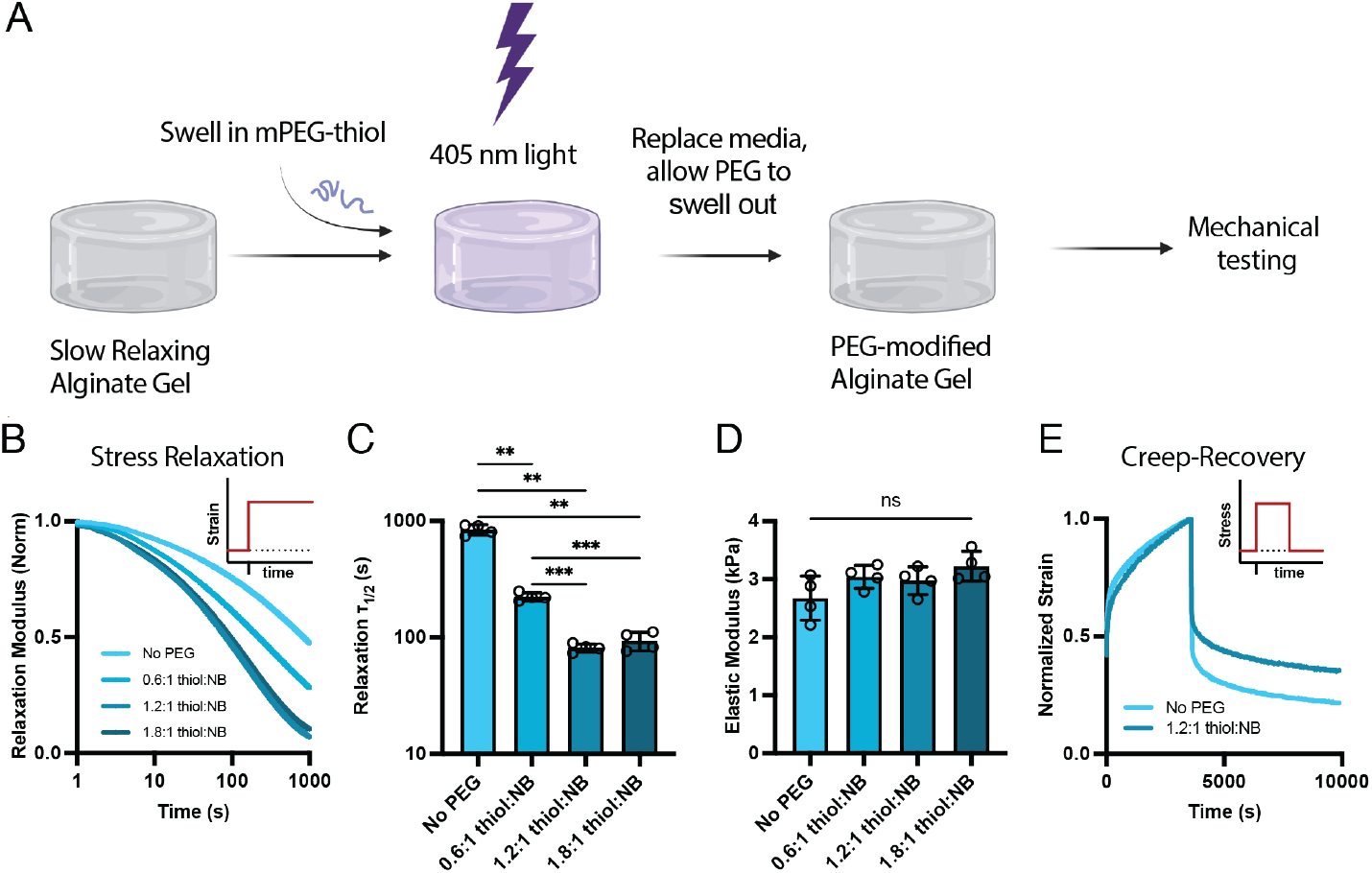
Light-triggered changes to alginate hydrogel viscoelasticity. (a) Schematic of experimental timeline. Norbornene-modified alginate hydrogels were equilibrated with a solution of mPEG-SH and LAP, exposed to 405 nm light, and unreacted PEG was allowed to diffuse out of the hydrogel before mechanical testing. (b,c) Stress relaxation rates can be enhanced by incorporating mPEG into the alginate network with a dependency on the amount of PEG added. (d) PEG photoconjugation does not significantly alter the hydrogel elastic modulus. (e) Creep-recovery tests demonstrate that PEG additional also alters the irrecoverable viscous deformation of the hydrogel network.

### Temporal increase in stress relaxation rate promotes cell spreading

After establishing our platform for phototunable viscoelastic properties, we then sought to determine how cells would respond to dynamic changes in stress relaxation rates. Alginate does not possess binding sites for cell adhesion, so peptides presenting the RGD-adhesion motif were coupled to norbornene-alginate to allow for cell adhesion^**11**^. Prior reports have demonstrated that mesenchymal stem cells (MSCs) have increasing protrusions and spreading in fast relaxing matrices compared to rounded morphologies in slow relaxing matrices^**11**^. We sought to determine if cell spreading could be induced on-demand by transitioning from slow relaxing to fast relaxing conditions in the presence of cells. (Fig. 3A). The amount of time in slow-relaxing conditions before PEG conjugation was varied for encapsulated MSCs. As expected, cells cultured in slow relaxing matrices for 7 days were highly spherical with few protrusions (Fig. 3B). Intriguingly, cells in matrices that were transitioned from slow relaxing to fast relaxing were significantly less spherical, had significantly larger volumes, and had more protrusions, captured by the solidity metric (Fig. 3B-E). These results show that the range of viscoelastic tunability of our approach is sufficient to observe differences in cellular response and that cells can respond to temporal changes to viscoelasticity. Interestingly, the magnitude of cell shape differences depended on the time of transition, with a diminished effect for cells transitioned later in the 7 day culture period (i.e. 3 slow/4 fast). Since both the time the cells were cultured in slow relaxing conditions and fast relaxing conditions were varied in this experimental design, the effects of each gel condition could not be decoupled.

**Figure 3:**
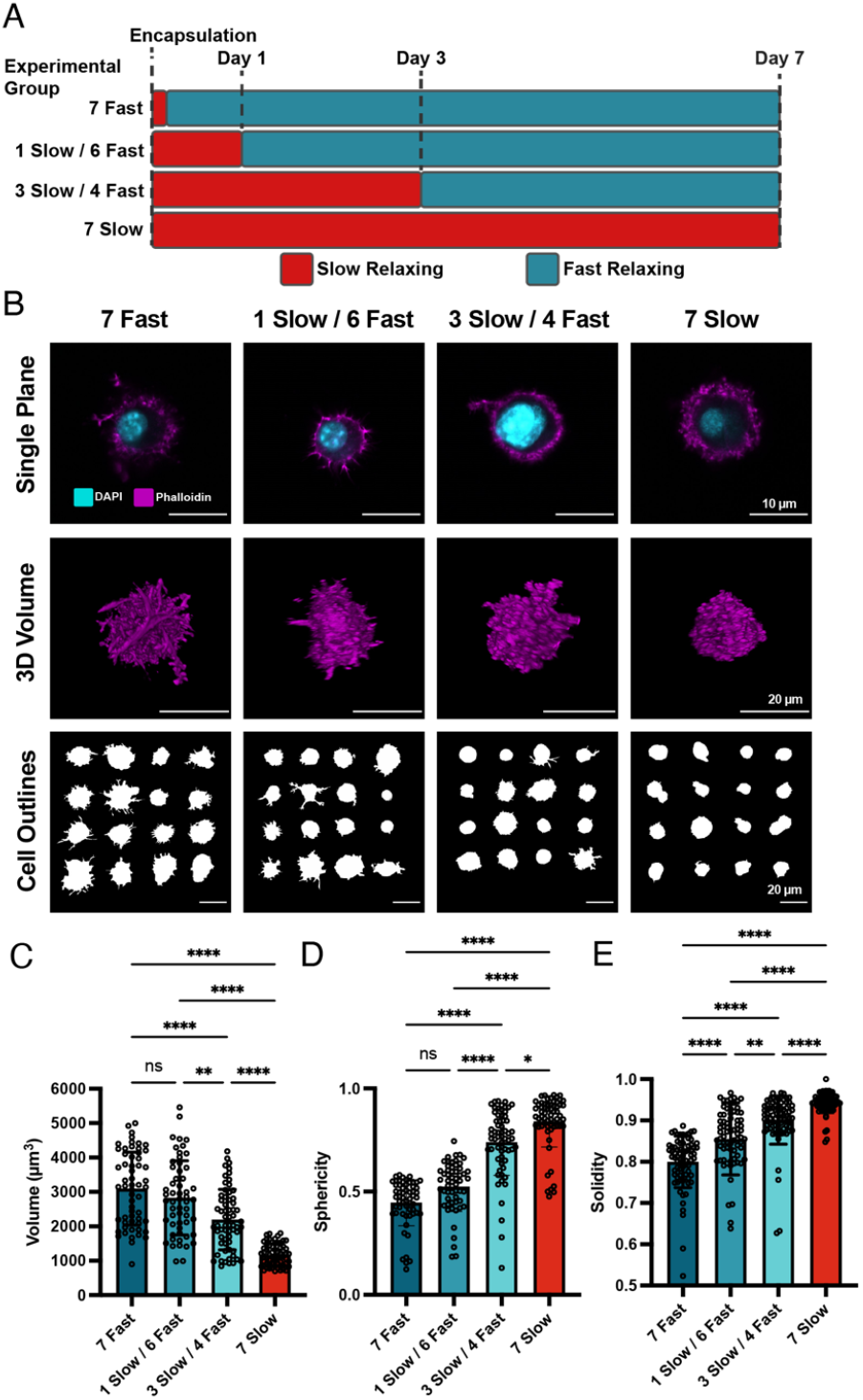
Cell spreading is responsive to dynamic viscoelasticity changes (a) Experimental timeline to evaluate cellular responses to changes in viscoelasticity on Day 0, 1, 3 or no changes (slow relaxing control). (b) MSCs exhibit spread morphologies when hydrogel stress relaxation is dynamically enhanced. Cross-sectional images of single MSCs, volumetric reconstructions of z-stacks, and outlines of maximum projections are shown. (c-e) Quantification of shape metrics (volume (c), sphericity (d), and solidity (e)) reveal significantly more spread and protrusive shapes as cells spend more time in fast relaxing hydrogels.

To distinguish the influence of the initial culture period in slow relaxing matrices from the total time in fast relaxing matrices, we varied the time in the initial slow relaxing matrices but maintained the cells in the fast-relaxing matrices for 7 days for all groups. (Fig. 4A). Between all groups cultured in fast relaxing conditions for 7 days, we did not observe any significant morphological differences, regardless of the initial time period in slow relaxing conditions (Fig. 7B). Regardless of the day of transition from slow to fast relaxing matrices, cells spreading was significantly different from cells in slow relaxing control gels. Cell volumes were similar between all groups in fast relaxing conditions, and higher than slow relaxing controls (Fig. 7C). Similarly, sphericity and solidity were significantly lower in each group that spent 7 days in fast relaxing conditions (Fig. 7 D-E). Together, these results indicate that MSCs morphology is responsive to dynamic matrix viscoelasticity. Additionally, cell spreading depends on the time spent in fast relaxing matrices and is not impeded by initial culture time in slow relaxing matrices, at least up to the 3 days in our experimental design.

**Figure 4:**
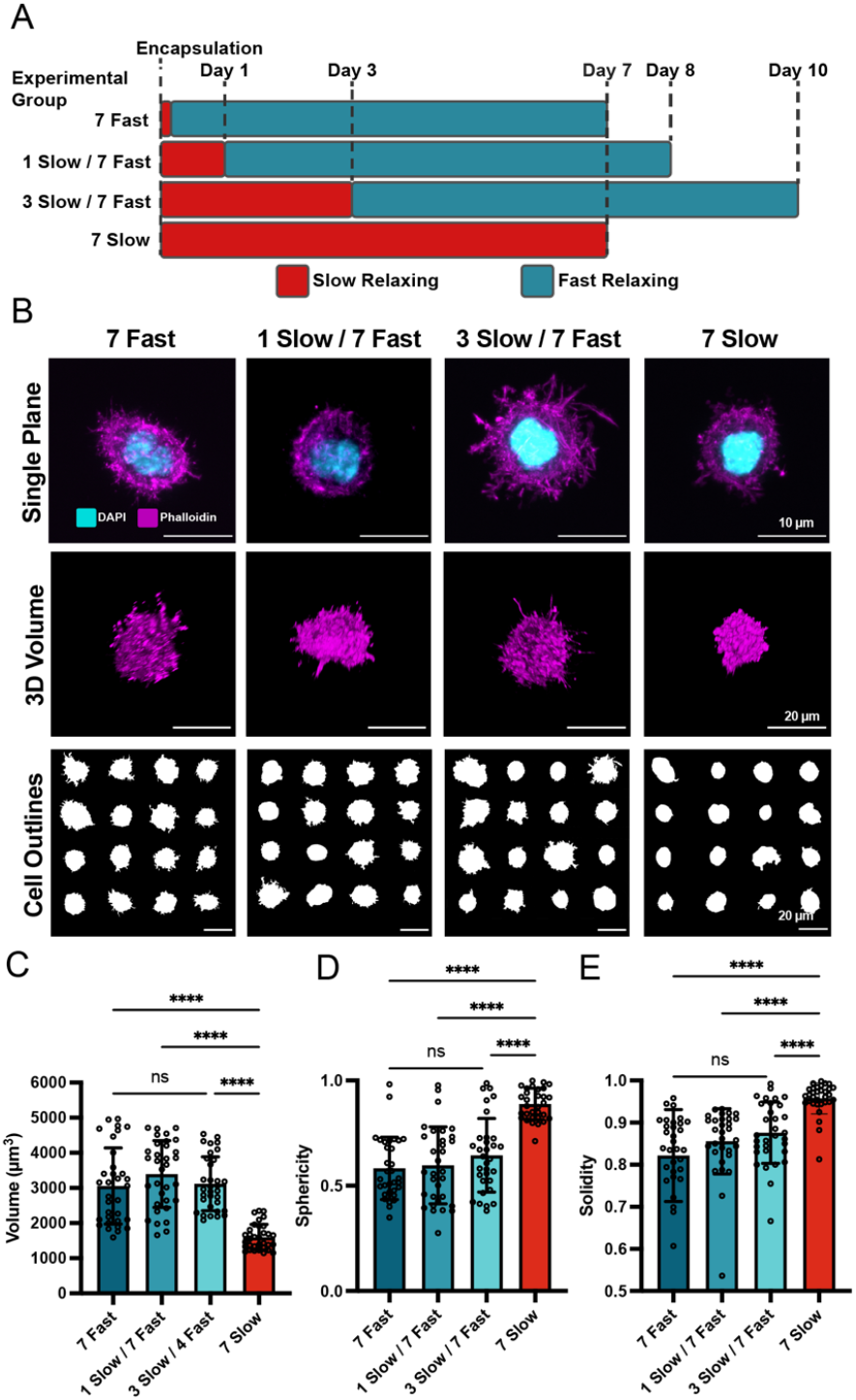
Spread morphologies depend on total time in fast relaxing conditions, not initial time in slow relaxing conditions. (a) Experimental timeline to evaluate morphological changes based on initial time in slow relaxing hydrogels. (b) MSCs exhibit spread morphologies after 7 days in fast relaxing hydrogels, regardless of initial time in slow relaxing conditions. Cross-sectional images of single MSCs, volumetric reconstructions of z-stacks, and outlines of maximum projections are shown. (c-e) Quantification of shape metrics (volume (c), sphericity (d), and solidity (e)) reveal cell spreading and protrusions do not significantly differ based on initial culture period in slow relaxing hydrogels.

### Transition from slow to fast relaxing matrices promotes proliferation

We next sought to evaluate the impact of dynamically altering the matrix stress relaxation rate on cell proliferation. Stress relaxation rate was recently shown to regulate proliferation and cell cycle progression in a metastatic breast cancer cell line, MDA-MB-231^**22**^. To determine if proliferation rate is also responsive to dynamic changes in matrix viscoelasticity, we cultured MDA-MB-231 cells in matrices that were transitioned from slow to fast relaxation rates at day 0, 1, or 3. All samples were fixed after 5 total days in culture (Fig. 5A). Cell proliferation was measured by incorporation EdU after 4 days of culture. Regardless of day of matrix transition, fast relaxing conditions produced significantly higher number of EdU-positive cells after 5 days compared to cells in slow relaxing conditions (Fig. 5B,C). There were no significant differences in the fraction of EdU-positive cells for matrices transitioned after 1 or 3 days compared to 5 days in fast relaxing only matrices.

**Figure 5:**
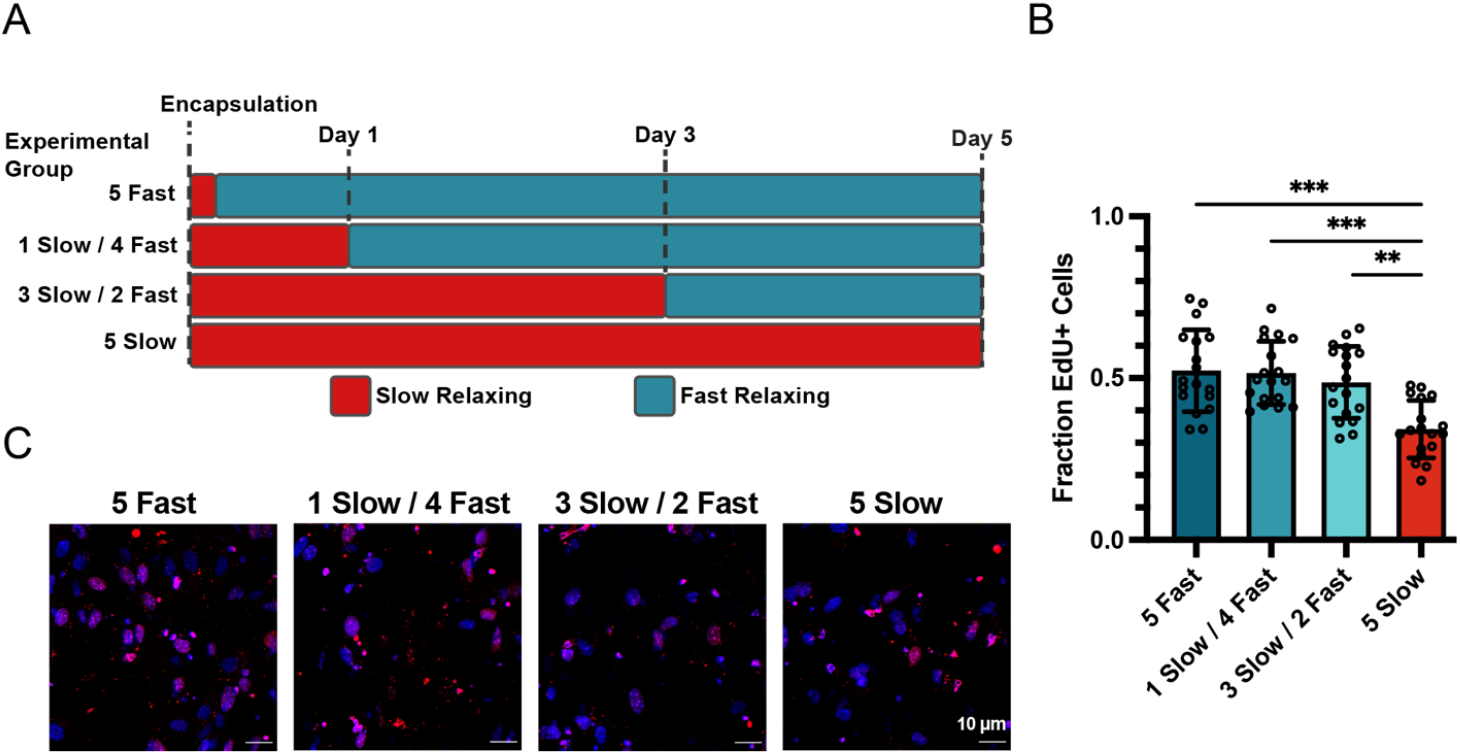
Proliferation rate increases when hydrogel viscoelasticity is dynamically enhanced. (a) Experimental timeline to evaluate proliferation after increasing the stress relaxation rate of hydrogels on Day 0, 1, 3 or in unchanged slow relaxing control conditions. (b) MDA-MB-231 cells are significantly more proliferative after dynamically increasing the stress relaxation rate of hydrogels compared to constant, slow relaxing controls. Proliferation rates do not significantly change based on the day of changing matrix viscoelasticity. EdU staining (red) indicates proliferative cells used for quantification in (b).

### Spatial patterning of viscoelasticity and effects on cell morphology

While photoaddition allows easy modification of alginate stress relaxation properties over time, our approach also enables photopatterning to spatial control of cellular behavior through matrix mechanics. To this end, we patterned hydrogels using a collimated laser source and a laser-printed photomask. To visualize the resulting patterns, 5% v/v of the PEG was labeled with FITC. Patterns could be easily made in a chambered cover glass or glass-bottomed well plate with good fidelity (Fig. 6A). To quantify the pattern fidelity in 3D with this system, we patterned the gel with lines of decreasing widths projected through the thickness of the gel (> 1 mm). Pattern fidelity in the x-y plane near the photomask was excellent (Fig. 6B,C). Deeper into the gel, 50 μm lines were preserved but 25 μm lines became distorted. Notably, the spatial resolution we achieved is still sufficient for all but patterning at the scale of single cells deep into hydrogels. Further, this limitation is a result of the optical setup, and could be overcome with more advanced photopatterning techniques that have been previously utilized with thiol-ene photochemistry^**48,49**^. To demonstrate the utility of this capability, we encapsulated MSCs in a uniformly slow relaxing 3D gel, then patterned 250 μm lines of fluorescent PEG to enhance stress relaxation. After 7 days of culture in the patterned gel, the cells were fixed and stained with phalloidin and DAPI. Using a tiled scan of whole gels to unbiasedly image the samples, different morphologies were observed in regions with and without PEG, indicating cellular responses to local viscoelasticity differences. (Fig. 6D). Cell morphologies in regions with and without PEG differed in area, circularity, and solidity (Fig. 6E-G). Cells in PEG patterned regions showed greater areas and decreased circularity and solidity, indicative of their greater number and size of protrusions. Overall, we demonstrate that this system can be used to pattern gel mechanics spatially, and that cells have a similar morphological response to being in a fast-relaxing local region of a gel as they do to being in an entirely fast-relaxing gel.

**Figure 6:**
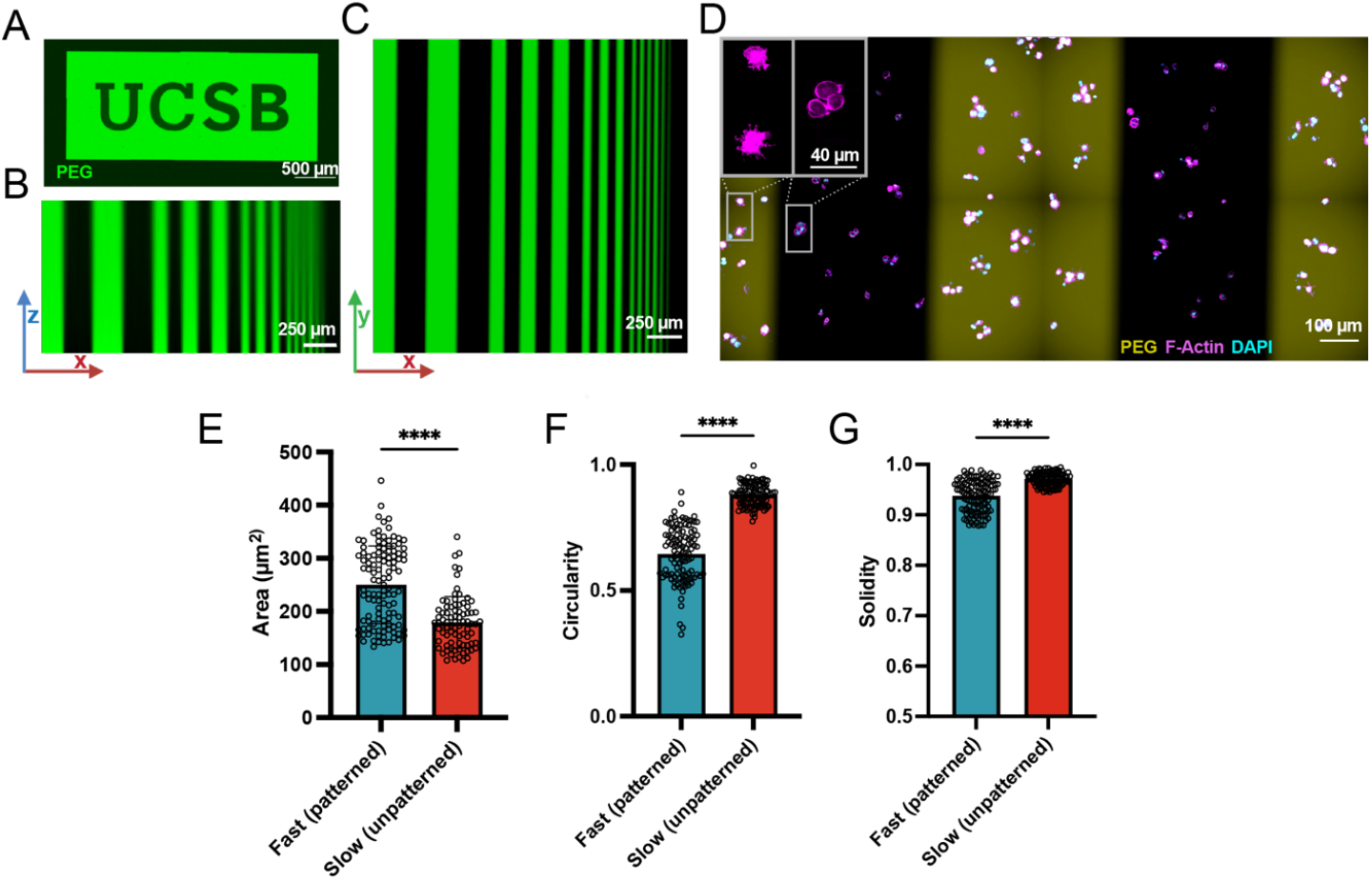
3D hydrogel stress relaxation rate can be spatially photopatterned. (a) Example photopattern to dynamically incorporate fluorescent PEGs to modulate stress relaxation rate locally. (b,c) Demonstration of spatial resolution via photomasking in the x-z plane (b) and x-y plane (c). Photopatterning of stress relaxation rates in the presence of cells (d). Cells in patterned regions of fast relaxation exhibit more spreading (e), are less rounded (f) and have more protrusions (g).

## CONCLUSIONS

We developed a method to increase the stress relaxation rate of 3D hydrogels in the presence of cells. Our light-triggered approach is able to modulate stress relaxation rates over the range in which cells sense and respond to matrix viscoelasticity. We found that cell protrusions, spreading, and shape are responsive to dynamic changes in matrix stress relaxation rate. Additionally, cell proliferation rate is also sensitive to changes in matrix viscoelasticity. We utilized the light-based approach to show high spatial control of 3D viscoelasticity by photopatterning as well, and again demonstrated the morphological response of cell in distinct viscoelastic environments. This platform addresses a critical unmet need for modeling spatiotemporally dynamic cellular microenvironments with 3D in vitro cell culture. We also envision this platform in applications for guiding or directing cell fate and tissue geometry over space and time.

## AUTHOR CONTRIBUTIONS

Philip Crandell performed the experiments and data analysis. Philip Crandell and Ryan Stowers conceptualized the experiments, wrote and edited the manuscript. Ryan Stowers is the principal investigator.

## ACKNOWLEDGEMENTS

We thank the UCSB Materials Research Laboratory for the use of the rheometer. The Materials Research Laboratory Shared Experimental Facilities are supported by the MRSEC Program of the NSF under Award No. DMR 1720256; a member of the NSF-funded Materials Research Facilities Network.

